# *Armillaria korhonenii*, the sixteenth biological species of *Armillaria* from China

**DOI:** 10.1101/2023.12.04.569844

**Authors:** Jianwei Liu, Guofu Qin, Jian Chen, Jing Song, Zhuyue Yan, Shimei Yang, Meng-hua Tian, Xin Xu, Changfei Zhang, Thatsanee Luangharn, Chitrabhanu S. Bhunjun, Fuqiang Yu, Zhuliang Yang

## Abstract

*Armillaria korhonenii*, designated as Chinese biological species Q (CBSQ) here, is described as a new species based on phylogenetic and morphological evidence as well as the results of mating tests. This species is the sixteenth biological species of *Armillaria* reported from China following the discovery of 15 previously known biological species. Phylogenetically, *A*. *korhonenii* is the earliest divergent lineage of the genus *Armillaria* as inferred from the nucleotide sequences of the Elongation Factor 1-Alpha (*EF1a*). Morphologically, it is characterized by its ellipsoid to elongate basidiospores, yellowish orange to honey yellow pileus with fibrillose squamules. Mating tests indicated that *A*. *korhonenii* is a tetrapolar heterogeneous species and is inter-sterile with all the tested species except partial compatibility with CBSH.

## INTRODUCTION

*Armillaria* (Fr.) Staude is a genus of Physalacriaceae, and the type species is *A. mellea* (Vahl: Fr.) P. Kumm (Kirk et al. 2008). Species belonging to *Armillaria* hold significant economic and ecological importance, with both positive and negative aspects from an anthropocentric perspective (Wu et al. 2019; Yuan et al. 2023). As serious pathogens, they can kill living plants, with detrimental consequences of forest destruction and substantial economic losses (Dai et al. 2007). On the other hand, as saprotrophs, they play a crucial role in carbon and mineral cycling within forest ecosystems (Heinzelmann et al. 2019). Furthermore, species of *Armillaria* can serve as symbionts of orchid plants and other fungi (Cha et al. 1995; Cha et al. 1996; Guo and Xu 1992; Guo et al. 2016; Kusano 1911; Kikuchi and Yamaji 2010; Terashita 1996; Xu and Mu 1990). Notably, the growth of *Gastrodia elata* Blume and *Polyporus umbellatus* (Pers.) Fr. (known as Tianma and Fuling respectively in the Traditional Chinese Medicine) relies on the assistance of *Armillaria* fungi.

To date the genus *Armillaria*, including the closely related genus *Desarmillaria* (Koch et al. 2017), comprises over 40 described species globally (Heinzelmann et al. 2019). In China, 16 Chinese biological species (CBS A to CBS P) of *Armillaria* were documented in the past (Qin et al. 2008; Zhao et al. 2008). During the establishment of the biological species CBSE, two initial strains were included. Subsequently, Zhao et al. (2000) modified and transferred these two strains to CBSC and CBSD, respectively. As a result, CBSE is currently vacant, and 15 CBS are left behind. Peng and Zhao (2020) described a new species, *A. xiaocaobaensis* J.H. Peng & C.L. Zhao based on morphological and phylogenetic analysis inferred from *EF1a* genes.

Over an extended period, defining species within *Armillaria* has proven challenging. Three operational species concepts, based on morphological, biological, and phylogenetic criteria respectively, have been employed to differentiate various taxa. However, these methods often yield conflicting results. For instance, Guo et al. (2016) revealed fourteen Chinese biological species of *Armillaria* could be assigned to 15 phylogenetic lineages. In most cases, each biological species aligned with one specific phylogenetic lineage. But in a few cases, a CBS was corresponded to two phylogenetic lineages, or two CBS were linked to one phylogenetic lineage. Liang et al. (2021) uncovered that biological species recognition did not accurately mirror the natural evolutionary relationships within *Armillaria*.

As the majority of presently recognized *Armillaria* species were initially described as biological species, a practice dating back to Korhonen (1978), and Anderson and Ullrich (1979) and given the limitations in resolving all species recognition issues with morphological and phylogenetic methods, our study prioritized the utilization of the biological species recognition method. A new *Armillaria* species with integrated biological, morphological and phylogenetic evidence is described herein.

## MATERIALS AND METHODS

### Biological studies.—

### Samples and spore prints collecting

Ten collections were freshly gathered from the Daweishan National Reserve and Ailaoshan National Reserve. After taking field photography and necessary fieldnotes, and isolating live strains, spore prints were prepared on sterile filter paper. These spore prints were then transported back to the laboratory for the isolation of individual spores, and the fungal fruitbodies were then dried at 45 C and deposited in the Herbarium of Cryptogams, Kunming Institute of Botany, Chinese Academy of Sciences (HKAS), Yunnan, China.

### The process of single-spore strains isolation and biological species determination

Spores obtained from a spore print were evenly distributed on the surface of water agar or a 1% malt-extract agar medium. The incubation period lasted for 1 to 3 days at a temperature of 24 C. Individual germinated spores were then selectively isolated using the technique outlined by Korhonen and Hintikka (1980). A minimum of 12 distinct single-spore cultures were obtained from each sample. The strains utilized for mating experiments are detailed in SUPPLEMENTARY TABLE 2. The procedures for mating experiments and determination of sexual compatibility followed the methods outlined by Zhao et al. (2008).

### Phylogenetic studies.—

### DNA extraction, PCR amplification and sequencing

DNA was extracted from dried specimens using CTAB method (Doyle 1987). To amplify *EF1α* gene sequences, we utilized universal primer pairs 983F/1567R (Rehner et al. 2001). The PCR procedure consisted of an initial pre-denaturation step at 94 C for 5 minutes, followed by 35 cycles of denaturation at 94 C for 40 seconds, annealing at 50 C for 40 seconds, and elongation at 72 C for 50 seconds. The process concluded with a final elongation step at 72 C for 8 minutes. Sequencing of the successful PCR products was carried out using 983F and 1567R primers on an ABI-3730-XL sequence analyzer from Applied Biosystems (Delaware).

### DNA sequence alignments and phylogenetic analyses

The *EF1α* sequences obtained from the specimens under this study were combined with those of the two studies available (Guo et al. 2016; Peng and Zhao 2020). Additionally, six sequences of *Desarmillaria ectypa* and eight of *D*. *tabescens* were incorporated as outgroup taxa, detailed information was shown in SUPPLEMENTARY TABLE 1. Subsequently, the dataset was aligned using MAFFT 6.8 (Katoh et al. 2005) and manually edited using BioEdit 7.0.9 as required (Hall 1999). Finally, the alignment of *EF1α* sequences was submitted to TreeBase (Study No. 30997).

Substitution model evaluation and the construction of a phylogenetic tree for maximum likelihood (ML) analysis were conducted using IQ-TREE 2.0.3 (Minh et al. 2020) on a Linux system. The substitution model chosen for the *EF1α* dataset was TNe+R4, determined through the Akaike information criterion (AIC) as suggested by IQ-TREE. Clade support in the ML analyses was assessed using the Shimodaira– Hasegawa-like approximate likelihood ratio test (SH-aLRT) with 1,000 replicates (Guindon et al. 2010), and 1,000 replicates of the ultrafast bootstrap (UFB) (Hoang et al. 2018). Nodes with support values of both SH-aLRT ≥ 80 and UFB ≥ 95 were considered well-supported, nodes with either SH-aLRT ≥ 80 or UFB ≥ 95 were considered weakly supported, and nodes with both SH-aLRT < 80 and UFB < 95 were deemed unsupported in the maximum likelihood (ML) analysis.

The *EF1α* dataset alignment consisted of 232 sequences, including 15 newly generated sequences, comprising 510 sites. Among these, 293 sites were conserved, and 275 were variable, with nearly 202 being parsimony-informative sites. The optimized log-likelihood value was -4665.462. Estimated base frequencies were as follows: A = 0.25, C = 0.25, G = 0.25, T = 0.25. Substitution rates were AC = 0.64487, AG = 3.29726, AT = 1.00000, CG = 0.64487, CT = 7.33867, GT = 1.00000, gamma distribution shape parameter α = 1.713.

### Morphological study.—

### Morphological observation

Dried specimens were meticulously hand-sectioned, mounted in 5% KOH, and observed using a Leica DM2500 compound microscope (Germany). Thirty basidiospores per collection were measured, and their sizes and shapes were documented and photographed. Color codes were recorded following Kornerup and Wanscher (1981). Basidiospore size was expressed as (a–) b – c (–d). Within the range b – c, at least 90% of the measured values were encompassed, while a and d denote the extremes of all measurements. Q signifies the length/width ratio of basidiospores in the side view, with Qm indicating the average Q of all basidiospores ± standard deviation. Descriptive terms adhere to the terminology established by Vellinga and Noordeloos (2001).

## RESULTS

### Biological studies.—

The pattern of sexuality of all haploid isolates from Daweishan National Reserve and Ailaoshan National Reserve showed a typical colony morphology of heterothallic sexual system. Thirty-six pairing between the haploid isolates representing eight specimens showed they belong to the same intersterility group (ISG). A total of 776 pairing combinations between the haploid isolates from Daweishan National Reserve and Ailaoshan National Reserve and the testers representing 15 Chinese Biological Species of *Armillaria*, from CBS A to P (except CBS E), and one European *A*. *cepistipes* and one North American *A*. *calvescens* have been carried out (SUPPLEMENTARY TABLE 2). As the result, the Daweishan National Reserve and Ailaoshan National Reserve’s ISGs were intersterile with all the tested species, except for partial compatible mating reaction between Daweishan National Reserve’s ISG and the CBS H. The mating test between 16 haploid isolates of CBS H and 8 haploid isolates of Daweishan National Reserve’s ISG demonstrated there was 8.3% compatible and 91.7% incompatible mating, respectively.

Therefore, the ISGs from Daweishan National Reserve and Ailaoshan National Reserve were identified as a new Chinese Biological Species and named as CBS Q in alphabetical order. All mating interactions among the species were summarized in the SUPPLEMENTARY TABLE 2.

### Phylogenetic studies.—

The phylogenetic tree derived from the *EF1a* dataset revealed that the ten collections formed a distinct clade, clustering with three collections of “*Armillaria* sp. 10” from Guo et al. (2016) (FIG. 1). This clade demonstrated robust support (SH-aLRT/UFB = 97.5/100) in Maximum Likelihood (ML) analyses and occupied a basal position within the genus *Armillaria*.

**Figure 1.**
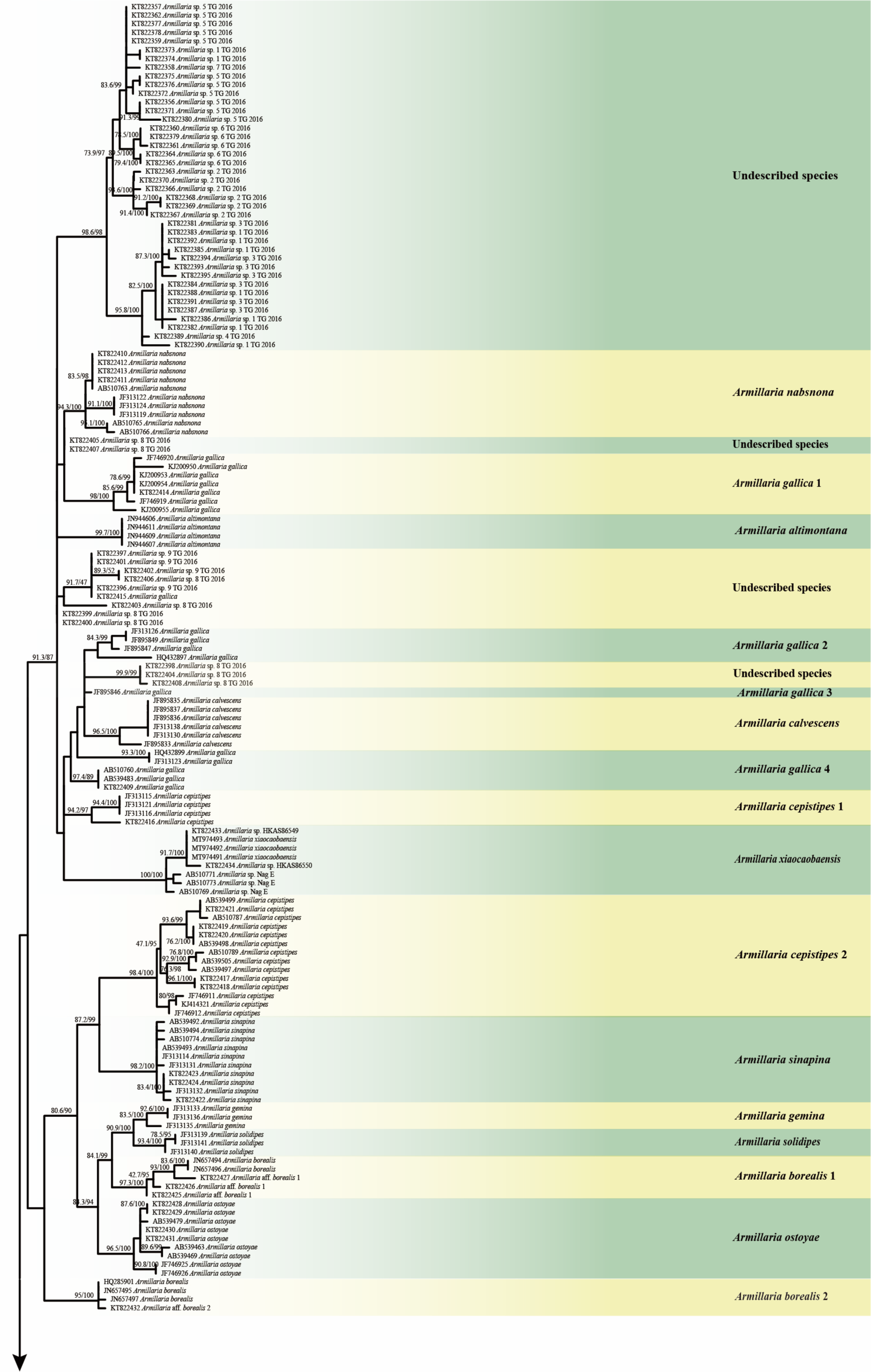
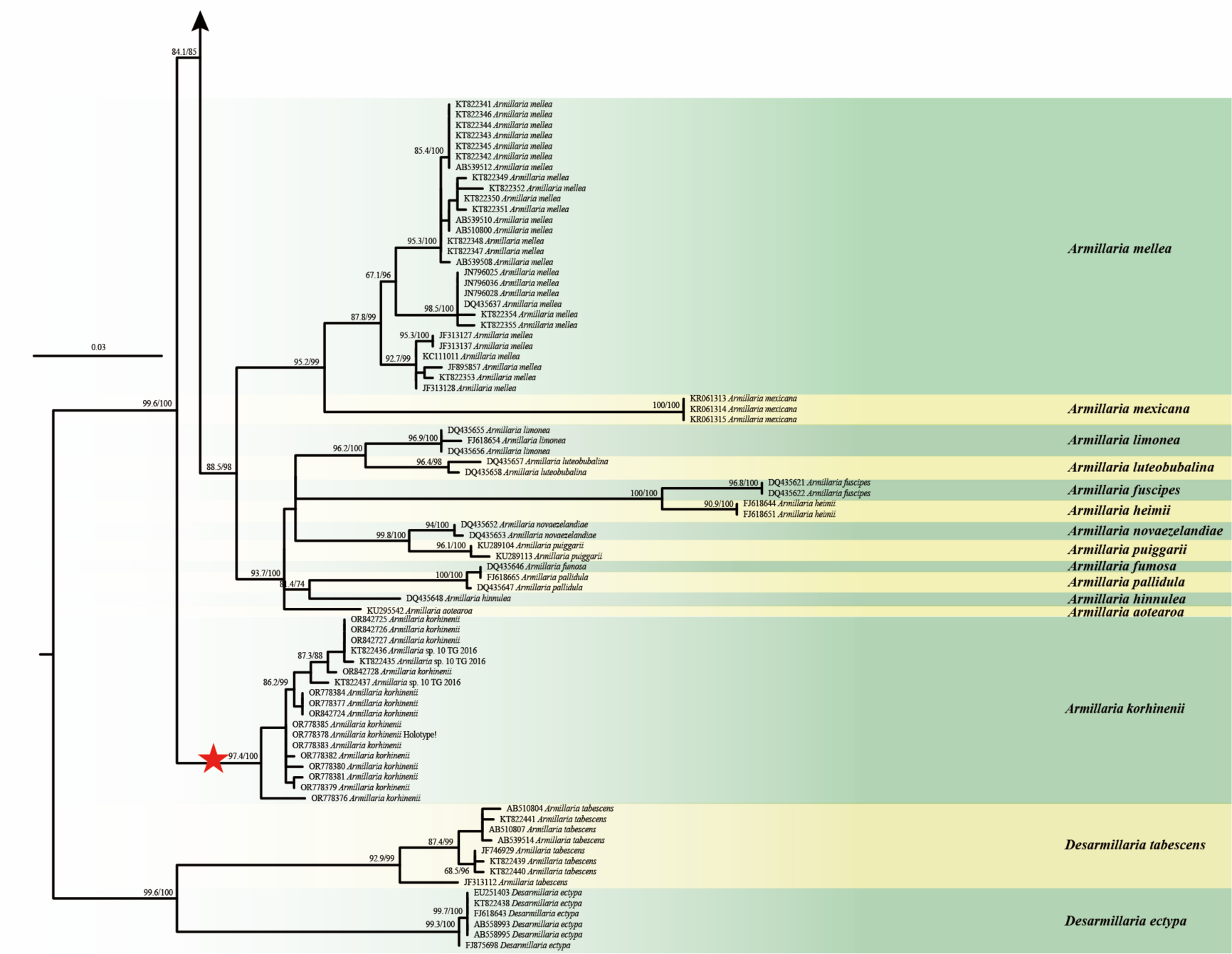
Maximum-Likelihood (ML) phylogenetic tree of *Armillaria* inferred from *EF1a* dataset, with Shimodaira–Hasegawa-like approximate likelihood ratio test (SH-aLRT) (left), ultrafast bootstrap (UFB) (right) near the corresponding node. Only one of SH-aLRT > 80 or UFB > 95 for ML was indicated along branches (SH-aLRT/UFB). *Armillaria korhonenii* is highlighted in bold and marked by a red asterisk.

## TAXONOMY

***Armillaria korhinenii*** J.W. Liu, G.F. Qin, J. Chen, F.Q. Yu & Zhu L. Yang, FIGS. 2–3

**Figure 2.**
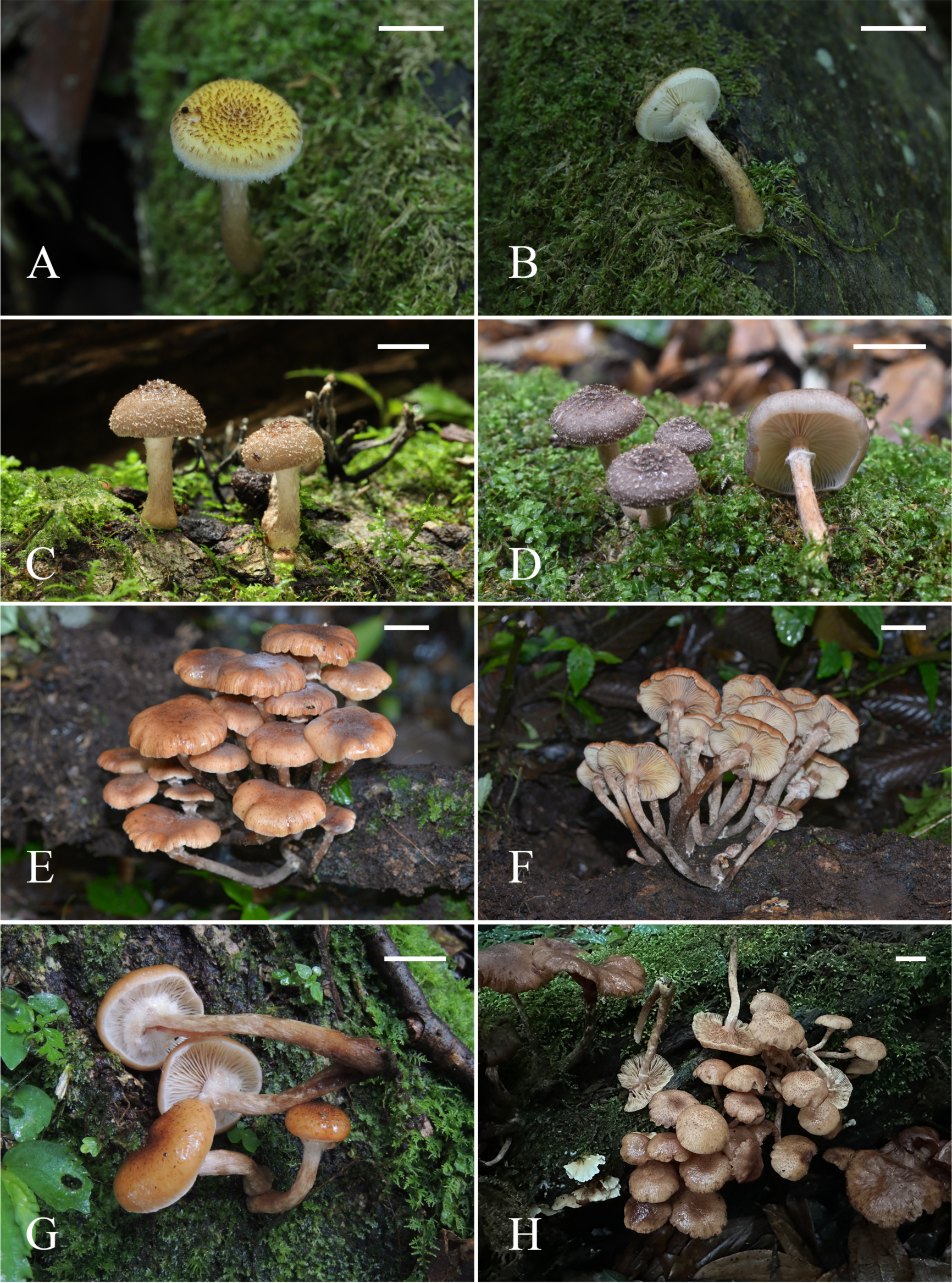
Basidiomata of *Armillaria korhonenii* at different stages of growth. A–B. WGS1417, photos by Gengshen Wang. C–D. WGS2045, Photos by Gengshen Wang. E–F. 22DWS03 (KUN-HKAS 126468), Photos by Jianwei Liu. G. 22DWS04 (KUN-HKAS 126469), Photo by Jianwei Liu. H. DAI 25212, Photo by Jian Chen. Bars: A–B = 1 cm, C = 1 cm, D = 1.5 cm, E–F = 3 cm, G = 1.5 cm, H = 1.5 cm.

**Figure 3.**
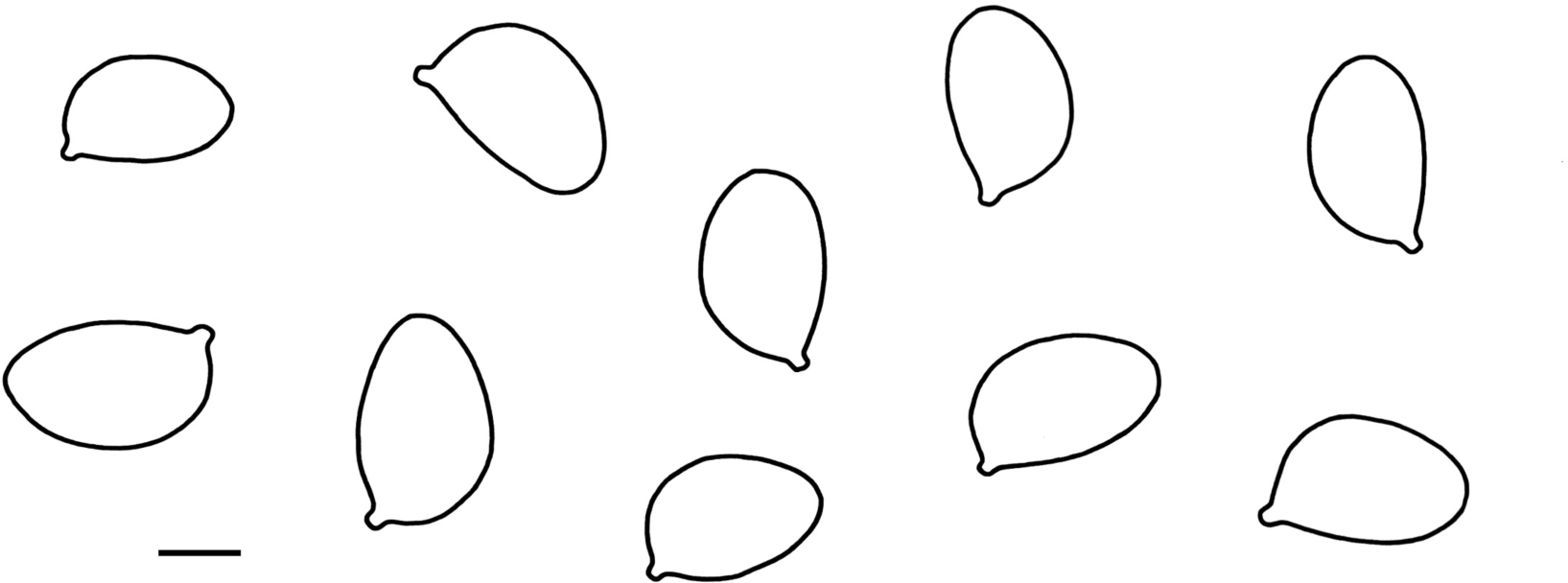
Basidiospores of *Armillaria korhonenii* (KUN-HKAS 126473, holotype). Drawings by Jianwei Liu. Bars = 4 μm.

MycoBank: MB850866

Typification: CHINA, YUNNAN: Pingbian County, Daweishan Natural Reserve, 22°54′42.99″N, 103°41′51.01″E, alt 2090 m, 25 May 2022, *22DWS08* (**holotype**, KUN-HKAS 126473, GenBank accession No.: *EF1a* = OR778378).

Diagnosis: Similar to the species of *Armillaria* with a membranous ring but differs in its densely repent or erect fibrillose squamules, unique basal lineage and incompatibility with all of the tested species.

Etymology: This species is named in honor of Prof. Kari Korhonen, in recognition of his exceptional four-decade-long dedication to mycology, particularly for his significant contributions to the establishment of the biological species concept and associated testing methods within the genus *Armillaria*.

Description: Basidiomata tricholomatoid, pileus sub-hemispherical, convex to plano-convex when young, plane to applanate when mature, up to 7 cm in diam.; surface yellowish orange (3A6–7), honey yellow (6B6–8, 6C6–8), grayish (1E1–2) to gray-black (1F1–2), densely covered with brown (5D5), taupe (5D5–8), light reddish brown (7B5–6), repent or erect fibrillose squamules when young, slightly sparse when mature; margin incurved, irregularly and fibrillose squamules derived from breaking up of the veil, white (1A1); context 0.5–3 mm thick. Lamellae decurrent to sub-decurrent, white (1A1), cream (1A1) when young, flesh color (7B4–6), slightly yellowish orange (5A5–6), to slightly yellowish (6B5–6). Stipe 50–90 × 5–9 mm, central, apex slightly striate, fibrous, clavate and often bulbous, honey yellow (6B6–8, 6C6–8), brown (6D6–7), dark gray (6F4–7) to dark brown (6F1–3), gradually darker from top to bottom when old; context white to whitish (1A1). Annulus present, cottony to membranous, white (1A1) to cream (1A1) when young, slightly brown (5D5) to light brown (7B4) when mature.

Basidiospores [60/2/2] 8–10 × 5–7 (7.5) μm, (Q = (1.2) 1.33–1.8, Qm = 1.51 ± 0.13), ellipsoid to elongate, colorless, hyaline, smooth, inamyloid. Basidia 25–30 × 8– 9 μm, clavate, 4-spored, often with a basal clamp connection; sterigmata 4–5 μm long. Pleurocystidia absent. Squamules on pileus surface comprised of sub-parallelly arranged hyphae 5–20 μm wide, with nearly colorless or yellowish, thin to slightly thick cell wall. Clamp connections common in all parts of basidioma.

Ecology/Substrate/Host: Single, scattered, or clustered on rotten wood of fagaceous planta in subtropical evergreen broad-leaved forests.

Distribution: Currently known from southern and middle Yunnan, southwestern China.

Other specimens examined: **CHINA. YUNNAN: Pingbian County, Daweishan Natural Reserve:** 22°55′13.40′′N, 103°41′51.78′′E, alt. 1945 m, 24 May 2022, *22DWS01* (KUN-HKAS 126466, GenBank accession no.: *EF1a* = OR778385); 22°57′50.44′′N, 103°42′6.17′′E, alt. 2109 m, 24 May 2022, *22DWS02* (KUN-HKAS 126467, GenBank accession no.: *EF1a* = OR778384); 22°54′21.91′′N, 103°41′51.58′′E, alt. 2124 m, 24 May 2022, *22DWS03* (KUN-HKAS 126468, GenBank accession no.: *EF1a* = OR778383); 22°54′27.52′′N, 103°41′52.14′′E, alt. 2099 m, 24 May 2022, *22DWS04* (KUN-HKAS 126469, GenBank accession no.: *EF1a* = OR778382); 22°54′40.11′′N, 103°42′0.71′′E, alt. 2073 m, 25 May 2022, *22DWS05* (KUN-HKAS 126470, GenBank accession no.: *EF1a* = OR778381); 22°54′45.83′′N, 103°41′58.12′′E, alt. 2072 m, 25 May 2022, *22DWS06* (KUN-HKAS 126471, GenBank accession no.: *EF1a* = OR778380); 22°54′48.32′′N, 103°41′57.95′′E, alt. 2086 m, 25 May 2022, *22DWS07* (KUN-HKAS 126472, GenBank accession no.: *EF1a* = OR778379); 22°55′50.30′′N, 103°41′20.72′′E, alt. 1858 m, 22 May 2021, *WGS1417* (GenBank accession no.: *EF1a* = OR778377).

**Jingdong County, Ailaoshan Natural Reserve:** 24°31′53.52′′N, 101°1′40.31′′E, alt. 2529 m, 4 Jul 2023, *WGS2045* (GenBank accession no.: *EF1a* = OR778376).

**Xinping County, Ailaoshan Natural Reserve:** 23°59′N, 101°30′E, alt. 2494 m, 29 Jun 2023, *Dai25210* (GenBank accession no.: *EF1a* = OR842724); 23°59′N, 101°30′E, alt. 2494 m, 29 Jun 2023, *Dai25211* (GenBank accession no.: *EF1a* = OR842725); 23°59′N, 101°30′E, alt. 2494 m, 29 Jun 2023, *Dai25212* (GenBank accession no.: *EF1a* = OR842726); 23°59′N, 101°30′E, alt. 2494 m, 29 Jun 2023, *Dai25213* (GenBank accession no.: *EF1a* = OR842727); 23°59′N, 101°30′E, alt. 2494 m, 29 Jun 2023, *Dai25214* (GenBank accession no.: *EF1a* = OR842728).

## DISCUSSION

The newly described species was confirmed as “*Armillaria* sp. 10” in Guo et al. (2016), as indicated in our phylogenetic analysis (FIG. 1). This suggested that the distribution of *A*. *korhonenii* can be expanded to western Yunnan, such as Lincang and Longling. *Armillaria korhonenii* is the sixteenth biological species of *Armillaria* from China following the discovery of 15 previously known biological species (CBSA, CBSB, CBSC, CBSD, CBSF, CBSG, CBSH, CBSI, CBSJ, CBSK, CBSL, CBSM, CBSN, CBSO and CBSP). Interestingly, the current phylogenetic tree constructed based on *EF1a* genes indicated that this new species might represent the earliest divergent lineage within the *Armillaria* genus. However, such inference should be verified using more extensive taxa sampling and more phylogenetic date in the future.

Despite the initial characterization of numerous *Armillaria* species as biological species, relying solely on biological species recognition presents significant limitations. Obtaining single spore strains for all identified species has proven to be a formidable task, and the assessment of sexcial compatibility and incompatibility often depended on the expertise of experienced researchers. In many instances, such as between CBS C and H, CBS A and CBS J, CBS G and CBS K, CBS H and CBS L, CBS H and CBS Q, as well as *A*. *cepistipes* and *A*. *sinapina*, signs of partial intersterility were recorded (Banik and Burdsall, 1998; Bérubé et al. 1996; Qin et al. 2007), adding complexity to the species recognition. Morphological species recognition faces challenges as well, given minimal differences in macro- and micro-characteristics among closely related *Armillaria* species (Antonin et al. 2009; Bérubé and Dessureault 1989; Park et al. 2018). Additionally, Steenwyk et al. (2023, 1) highlighted that “biological factors, such as incomplete lineage sorting, horizontal gene transfer, hybridization, introgression, recombination and convergent molecular evolution, can lead to gene phylogenies that differ from the species tree”. Therefore, relying solely on a single or multi-gene phylogenetic analysis may not fully capture the true evolutionary history and relationships among different species.

Adopting a polyphasic approach becomes essential, integrating genetic lineages with considerations of biological (reproductive), morphological, and functional distinctions (Heinzelmann et al. 2019). As genomic data becomes more accessible, coupled with advanced analytical tools, our comprehension of the evolutionary history and inter-species relationships within the *Armillaria* genus is poised for significant advancement. Utilizing this well-established evolutionary history, the formulation of a comprehensive species classification concept, supported by a variety of evidence, holds the potential to effectively address historical classification challenges associated with the *Armillaria* genus.

## ACKNOWLEDGMENTS

We sincerely thank Mr. Gengshen Wang from Kunming Institute of Botany, Chinese Academy of Sciences for providing precise collections and images, and Prof. Yucheng Dai from School of Ecology and Nature Conservation, Beijing Forestry University for revising early manuscript.

---

**Supplementary Table 1.**
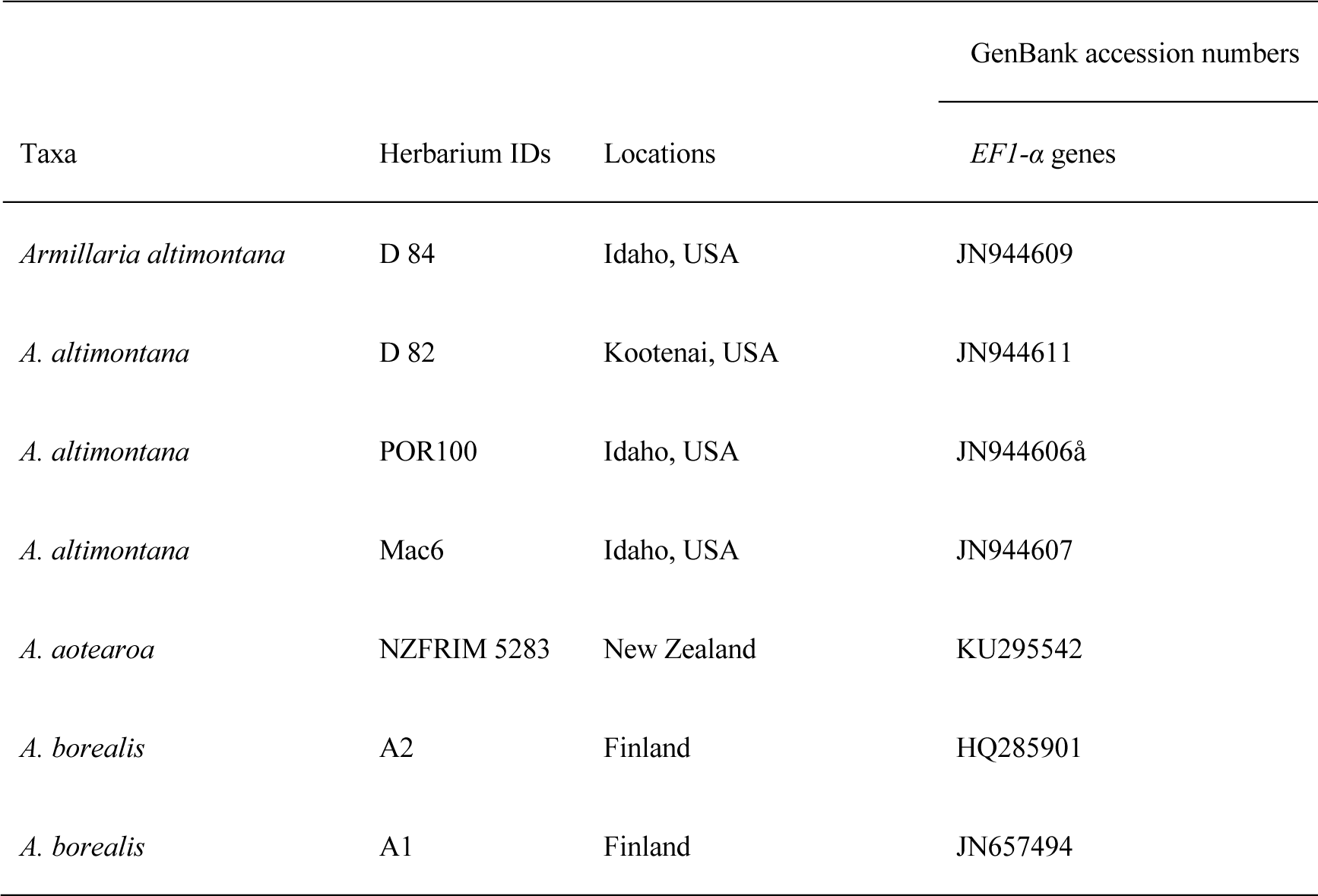
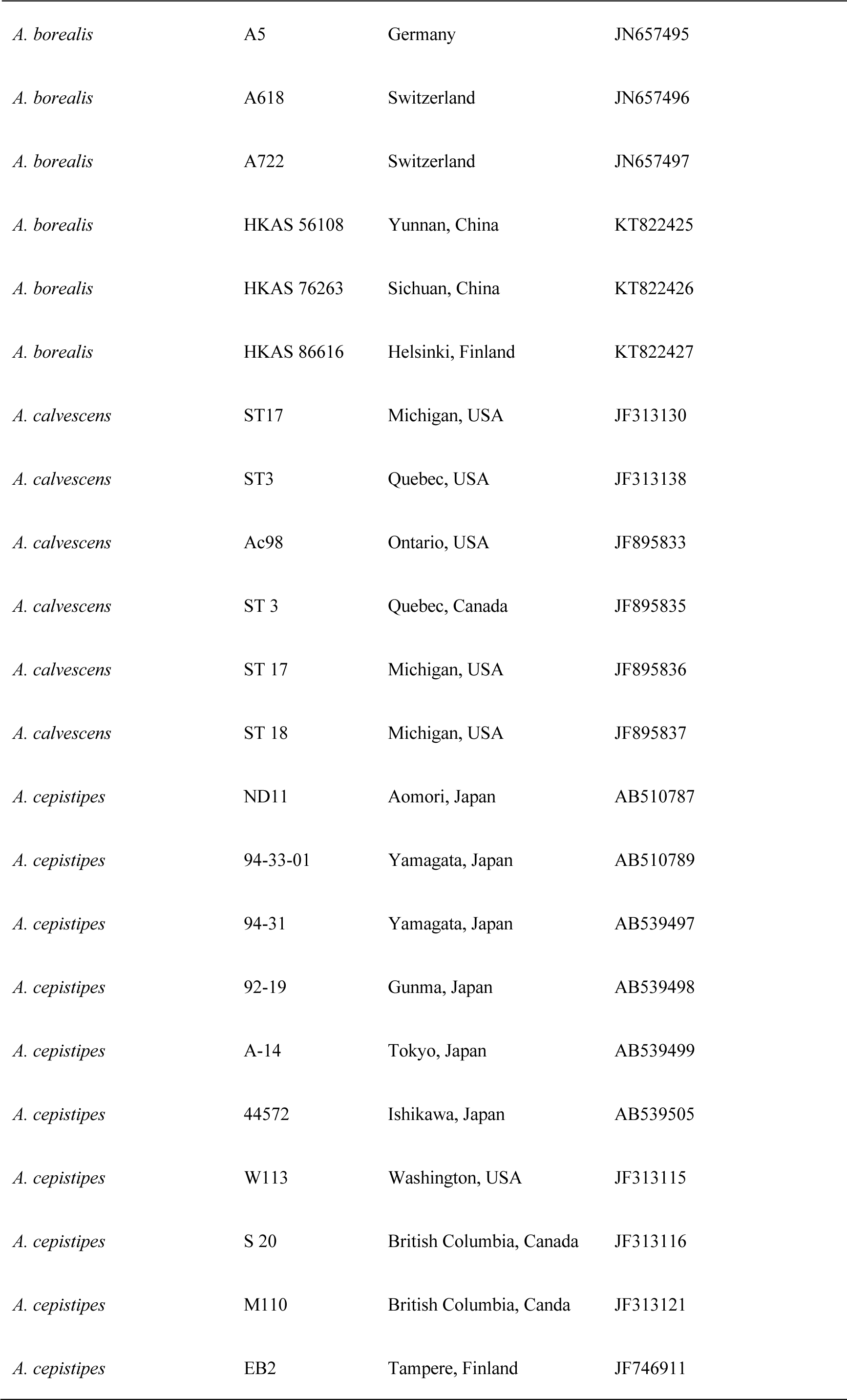
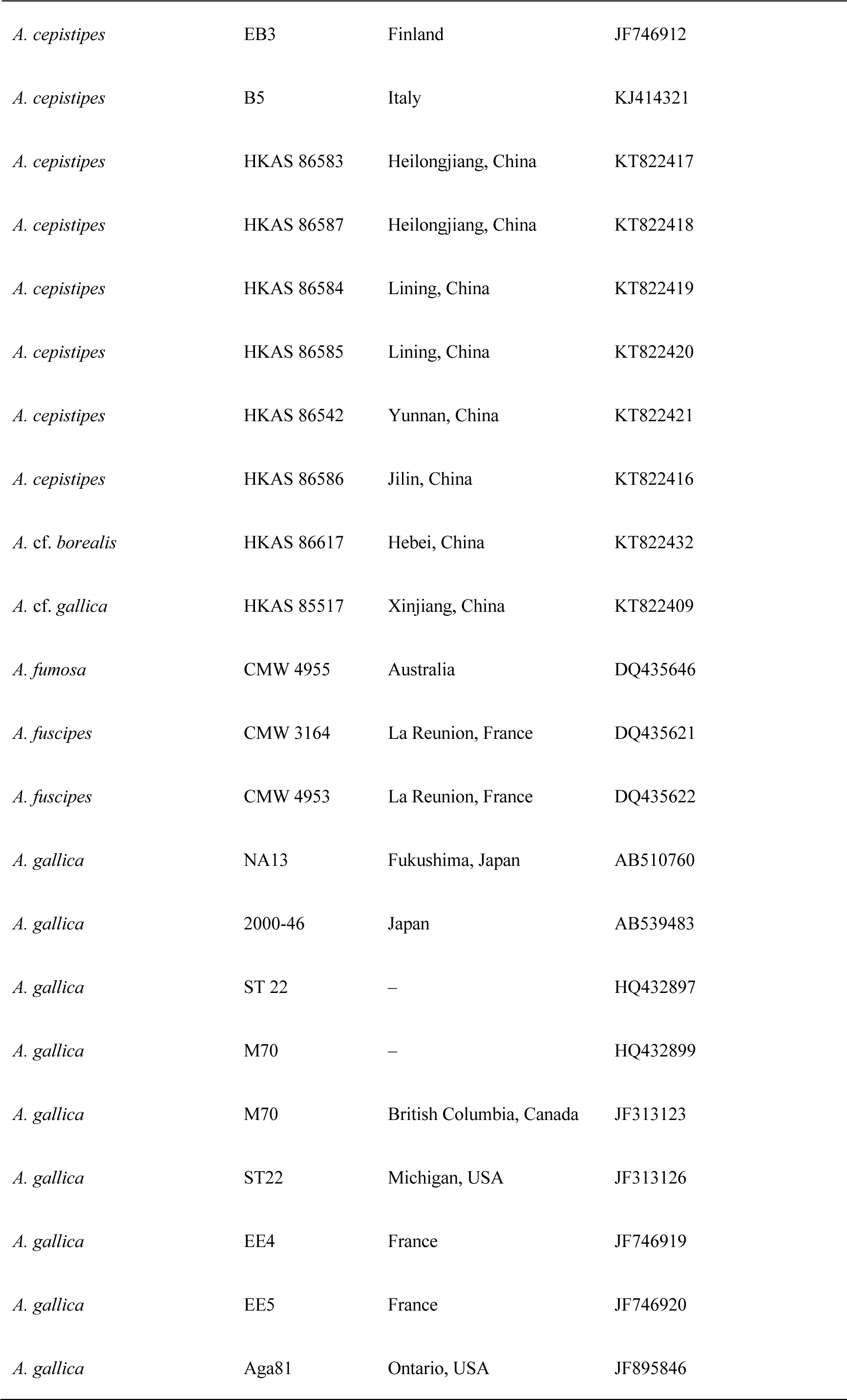
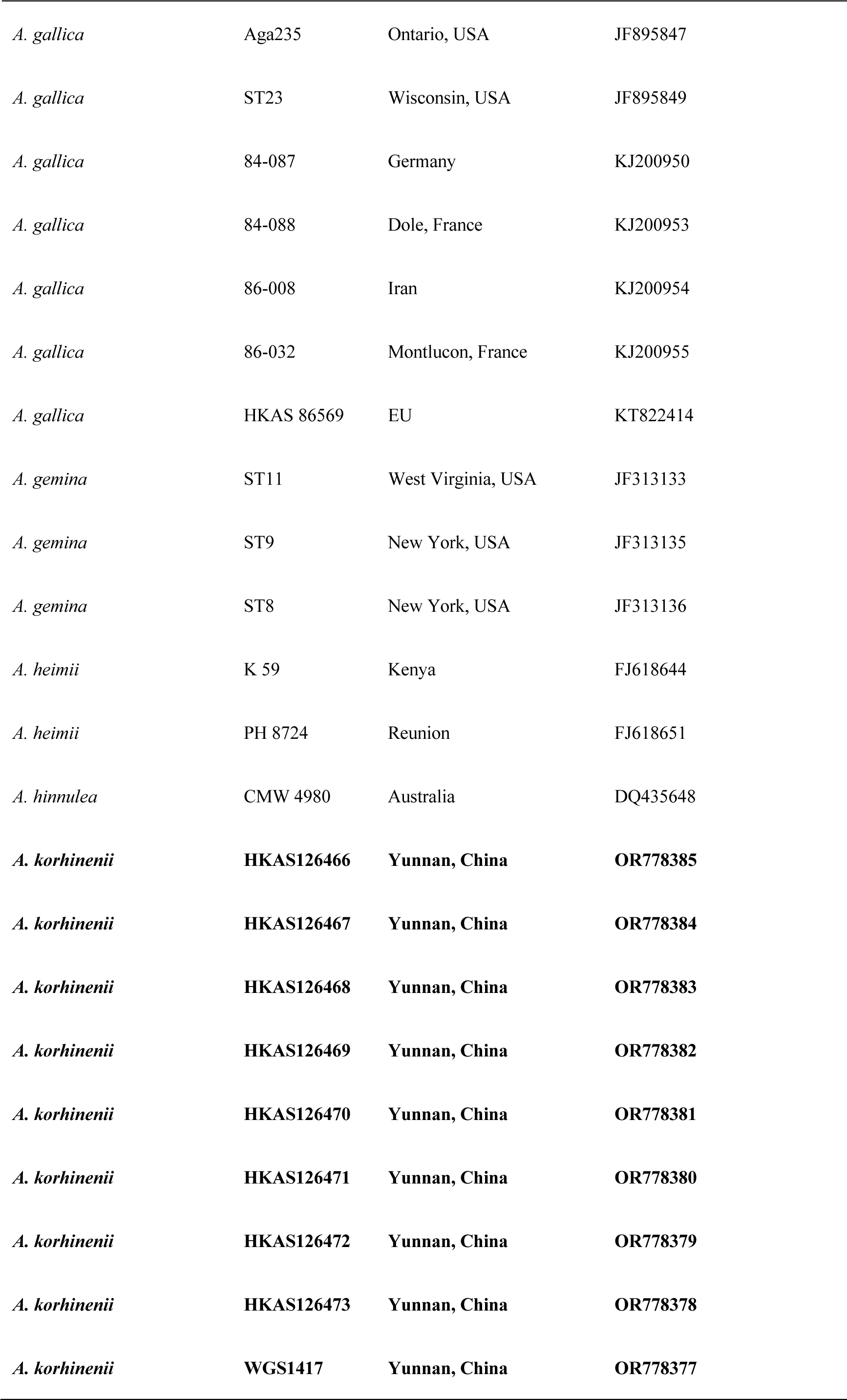
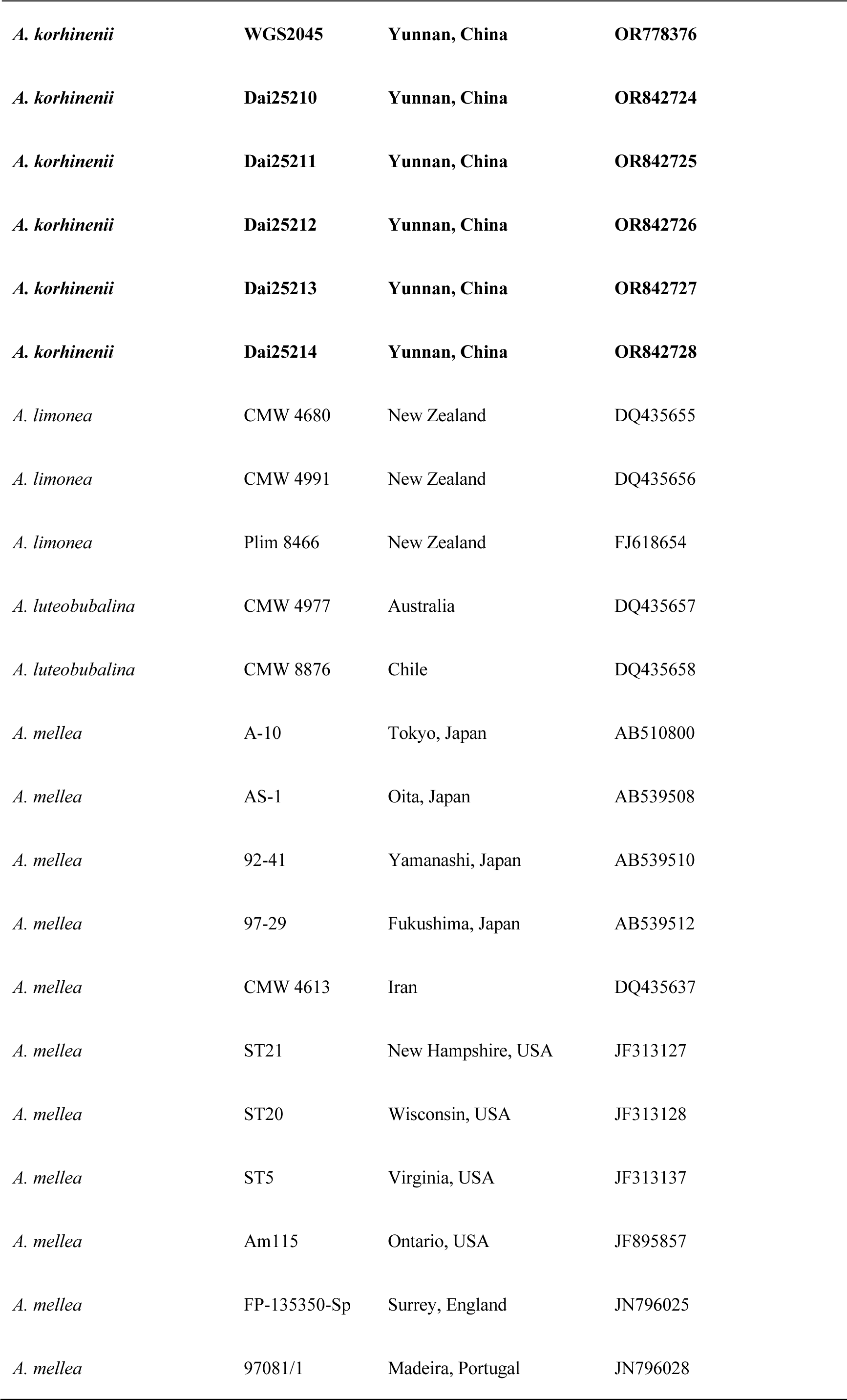
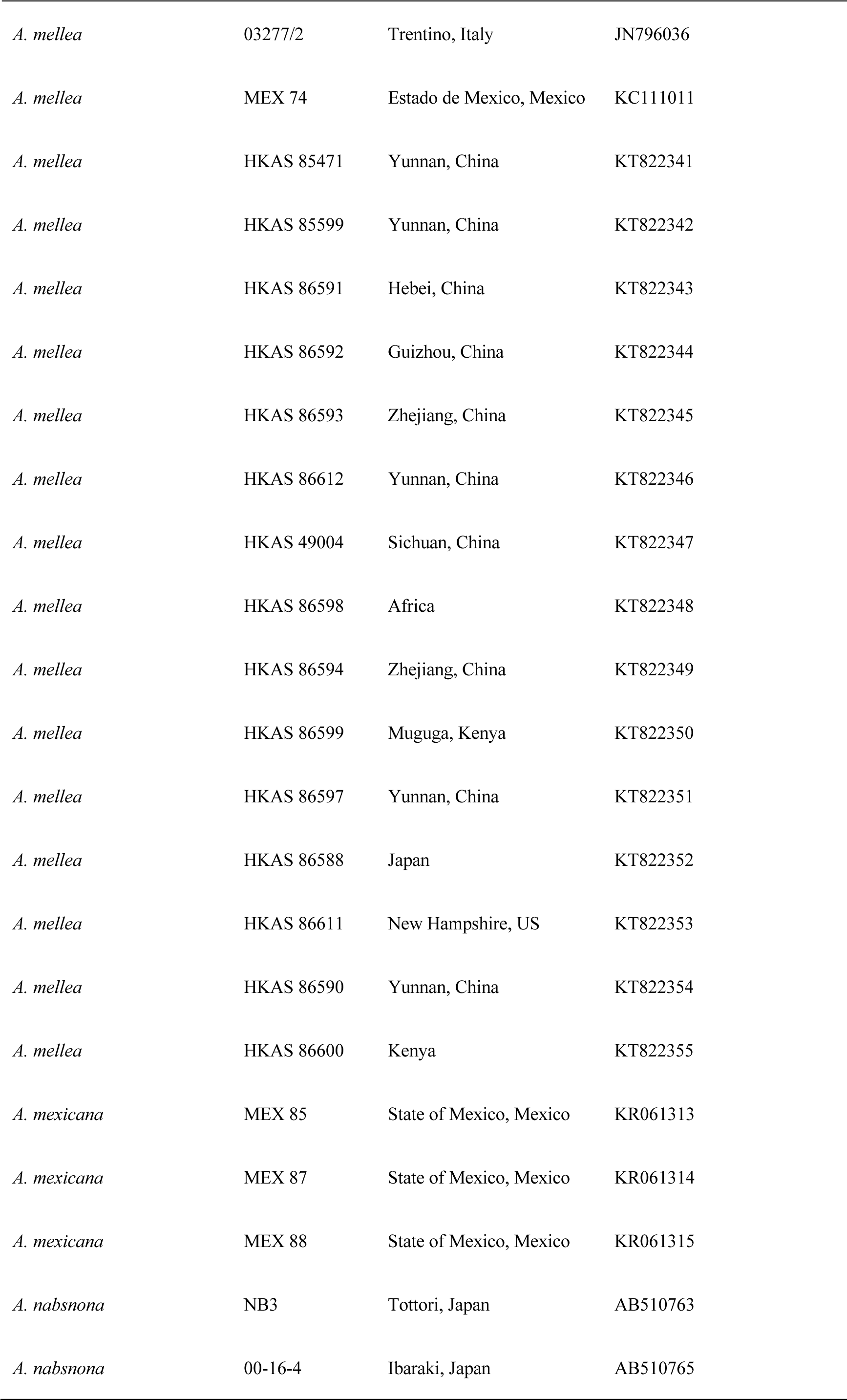
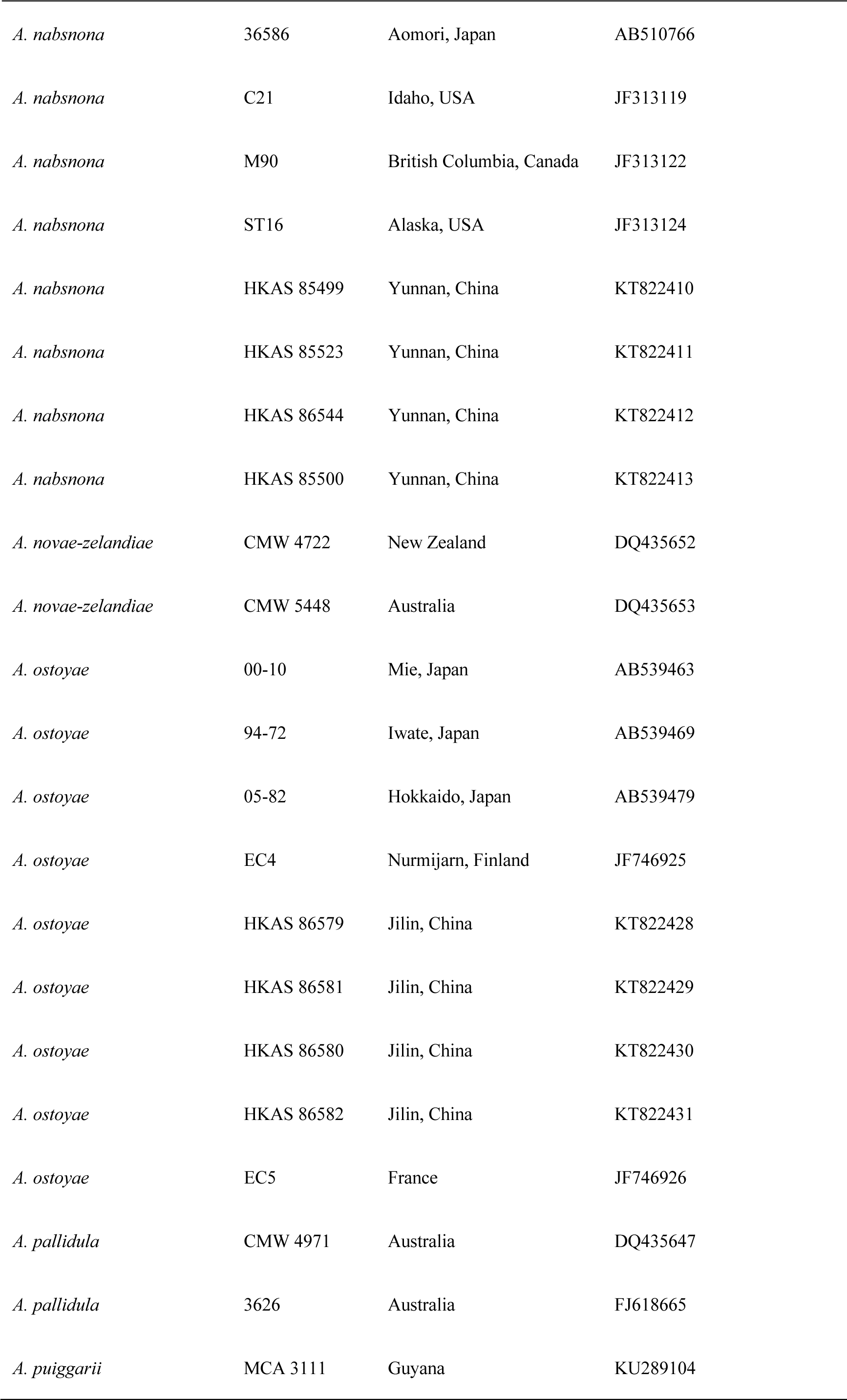
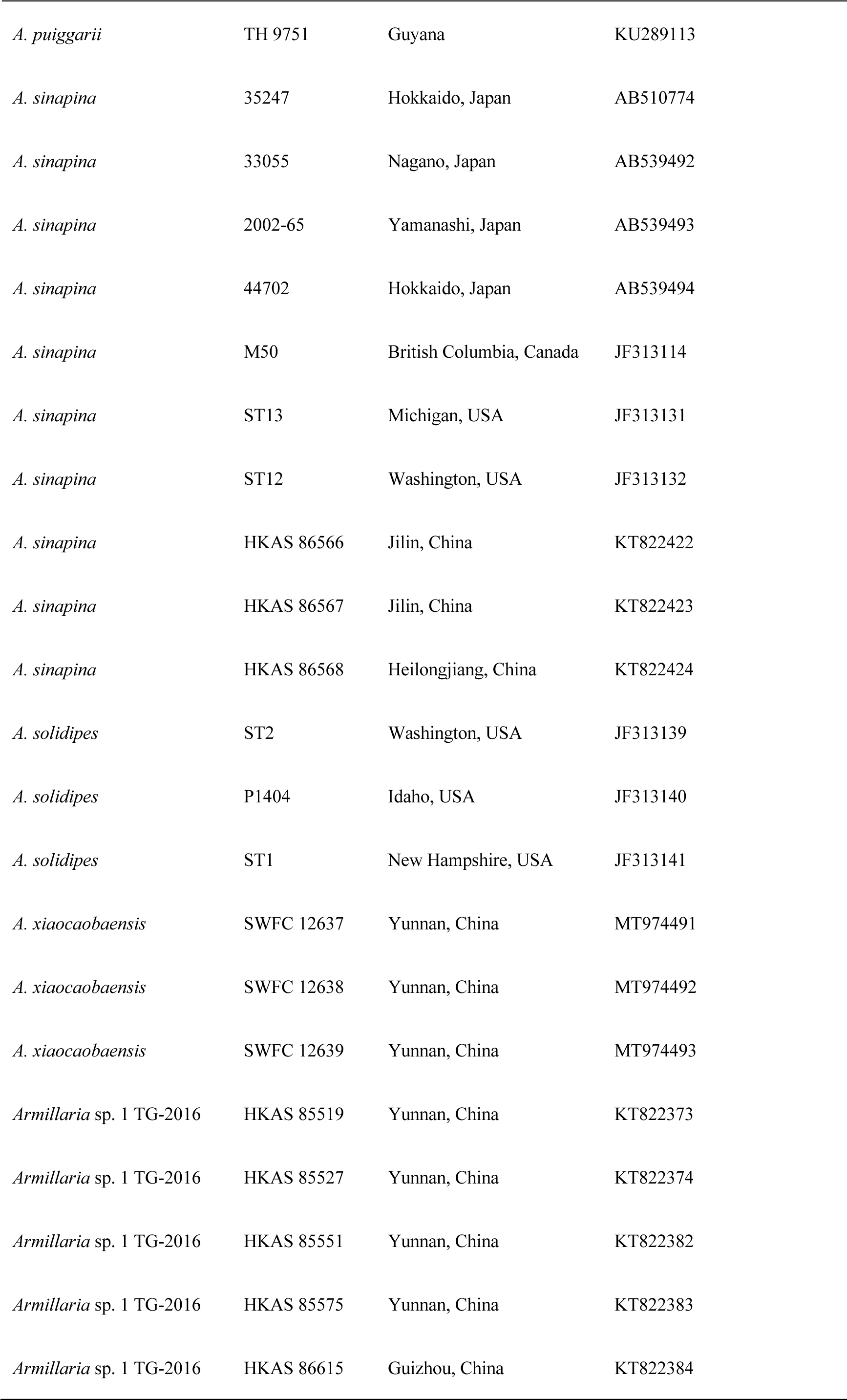
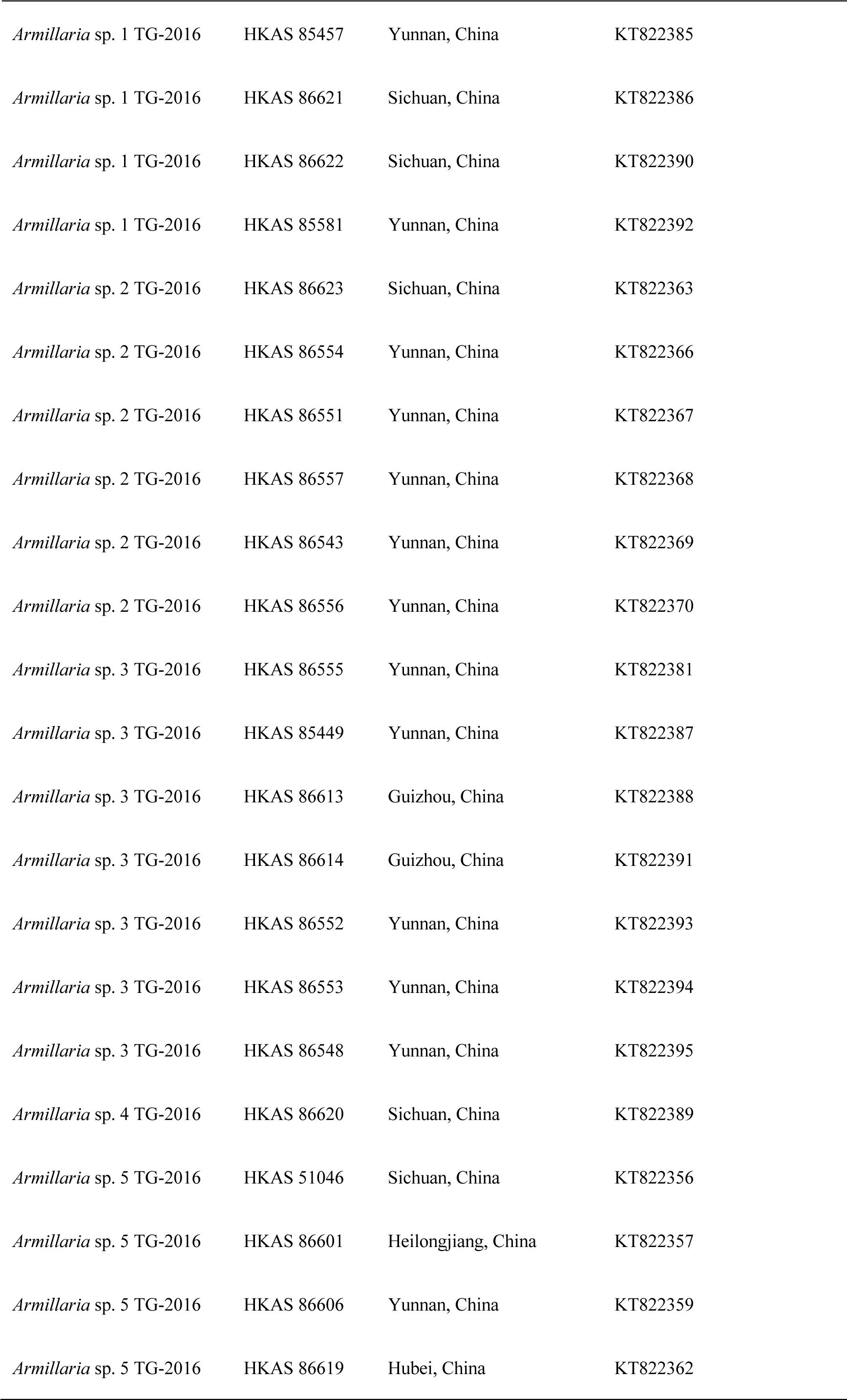
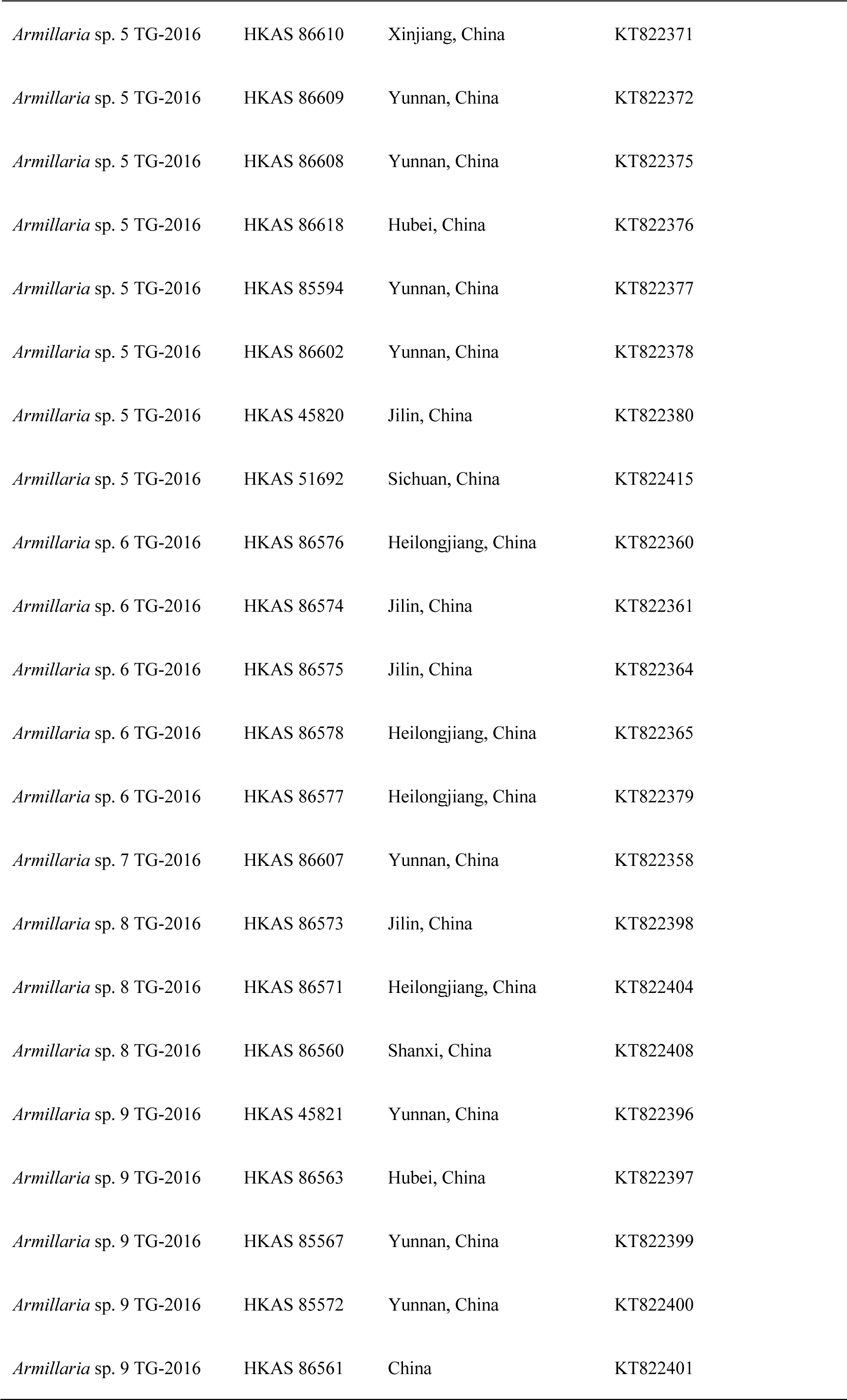
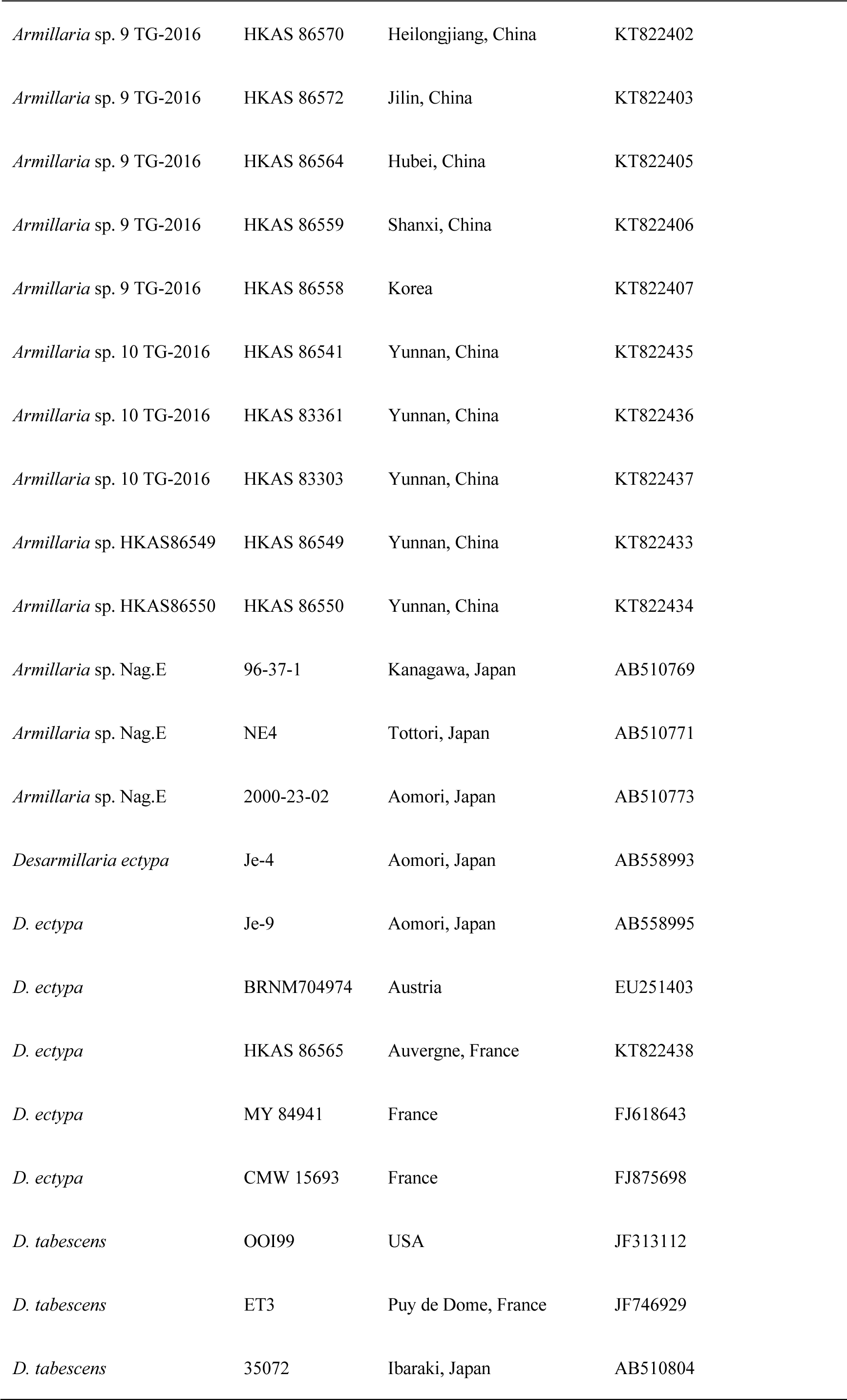
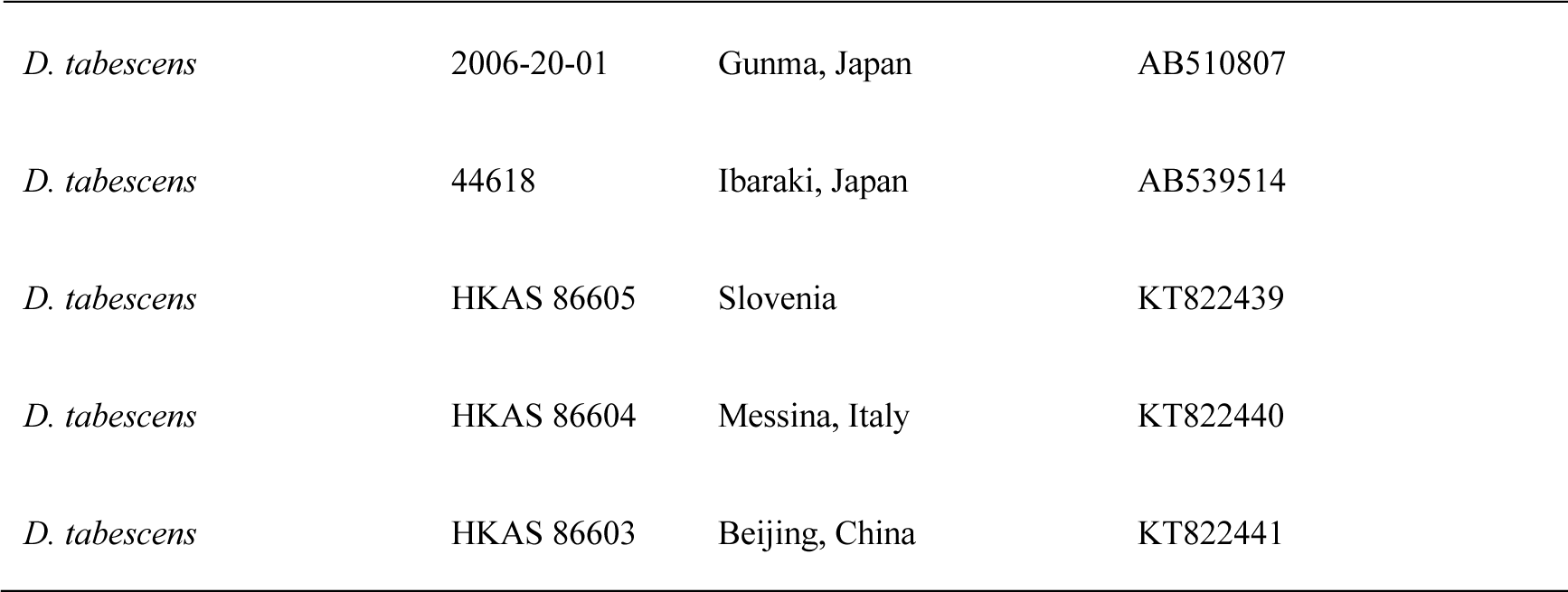
The names, voucher numbers, locations, and corresponding GenBank numbers of the taxa utilized in the phylogenetic analyses of the present study, with newly generated sequences highlighted in bold.

| Taxa | Herbarium IDs | Locations | GenBank accession numbers |
| --- | --- | --- | --- |
|  |  |  | *EF1-α* genes |
| *Armillaria altimontana* | D 84 | Idaho, USA | JN944609 |
| *A. altimontana* | D 82 | Kootenai, USA | JN944611 |
| *A. altimontana* | POR100 | Idaho, USA | JN944606 |
| *A. altimontana* | Mac6 | Idaho, USA | JN944607 |
| *A. aotearoa* | NZFRIM 5283 | New Zealand | KU295542 |
| *A. borealis* | A2 | Finland | HQ285901 |
| *A. borealis* | A1 | Finland | JN657494 |
| *A. borealis* | A5 | Germany | JN657495 |
| *A. borealis* | A618 | Switzerland | JN657496 |
| *A. borealis* | A722 | Switzerland | JN657497 |
| *A. borealis* | HKAS 56108 | Yunnan, China | KT822425 |
| *A. borealis* | HKAS 76263 | Sichuan, China | KT822426 |
| *A. borealis* | HKAS 86616 | Helsinki, Finland | KT822427 |
| *A. calvescens* | ST17 | Michigan, USA | JF313130 |
| *A. calvescens* | ST3 | Quebec, USA | JF313138 |
| *A. calvescens* | Ac98 | Ontario, USA | JF895833 |
| *A. calvescens* | ST 3 | Quebec, Canada | JF895835 |
| *A. calvescens* | ST 17 | Michigan, USA | JF895836 |
| *A. calvescens* | ST 18 | Michigan, USA | JF895837 |
| *A. cepistipes* | ND11 | Aomori, Japan | AB510787 |
| *A. cepistipes* | 94-33-01 | Yamagata, Japan | AB510789 |
| *A. cepistipes* | 94-31 | Yamagata, Japan | AB539497 |
| *A. cepistipes* | 92-19 | Gunma, Japan | AB539498 |
| *A. cepistipes* | A-14 | Tokyo, Japan | AB539499 |
| *A. cepistipes* | 44572 | Ishikawa, Japan | AB539505 |
| *A. cepistipes* | W113 | Washington, USA | JF313115 |
| *A. cepistipes* | S 20 | British Columbia, Canada | JF313116 |
| *A. cepistipes* | M110 | British Columbia, Canda | JF313121 |
| *A. cepistipes* | EB2 | Tampere, Finland | JF746911 |
| *A. cepistipes* | EB3 | Finland | JF746912 |
| *A. cepistipes* | B5 | Italy | KJ414321 |
| *A. cepistipes* | HKAS 86583 | Heilongjiang, China | KT822417 |
| *A. cepistipes* | HKAS 86587 | Heilongjiang, China | KT822418 |
| *A. cepistipes* | HKAS 86584 | Lining, China | KT822419 |
| *A. cepistipes* | HKAS 86585 | Lining, China | KT822420 |
| *A. cepistipes* | HKAS 86542 | Yunnan, China | KT822421 |
| *A. cepistipes* | HKAS 86586 | Jilin, China | KT822416 |
| *A. cf. borealis* | HKAS 86617 | Hebei, China | KT822432 |
| *A. cf. gallica* | HKAS 85517 | Xinjiang, China | KT822409 |
| *A. fumosa* | CMW 4955 | Australia | DQ435646 |
| *A. fuscipes* | CMW 3164 | La Reunion, France | DQ435621 |
| *A. fuscipes* | CMW 4953 | La Reunion, France | DQ435622 |
| *A. gallica* | NA13 | Fukushima, Japan | AB510760 |
| *A. gallica* | 2000-46 | Japan | AB539483 |
| *A. gallica* | ST 22 | — | HQ432897 |
| *A. gallica* | M70 | — | HQ432899 |
| *A. gallica* | M70 | British Columbia, Canada | JF313123 |
| *A. gallica* | ST22 | Michigan, USA | JF313126 |
| *A. gallica* | EE4 | France | JF746919 |
| *A. gallica* | EE5 | France | JF746920 |
| *A. gallica* | Aga81 | Ontario, USA | JF895846 |
| *A. gallica* | Aga235 | Ontario, USA | JF895847 |
| *A. gallica* | ST23 | Wisconsin, USA | JF895849 |
| *A. gallica* | 84-087 | Germany | KJ200950 |
| *A. gallica* | 84-088 | Dole, France | KJ200953 |
| *A. gallica* | 86-008 | Iran | KJ200954 |
| *A. gallica* | 86-032 | Montlucon, France | KJ200955 |
| *A. gallica* | HKAS 86569 | EU | KT822414 |
| *A. gemina* | ST11 | West Virginia, USA | JF313133 |
| *A. gemina* | ST9 | New York, USA | JF313135 |
| *A. gemina* | ST8 | New York, USA | JF313136 |
| *A. heimii* | K 59 | Kenya | FJ618644 |
| *A. heimii* | PH 8724 | Reunion | FJ618651 |
| *A. hinnulea* | CMW 4980 | Australia | DQ435648 |
| *A. korhinenii* | **HKAS126466** | **Yunnan, China** | **OR778385** |
| *A. korhinenii* | **HKAS126467** | **Yunnan, China** | **OR778384** |
| *A. korhinenii* | **HKAS126468** | **Yunnan, China** | **OR778383** |
| *A. korhinenii* | **HKAS126469** | **Yunnan, China** | **OR778382** |
| *A. korhinenii* | **HKAS126470** | **Yunnan, China** | **OR778381** |
| *A. korhinenii* | **HKAS126471** | **Yunnan, China** | **OR778380** |
| *A. korhinenii* | **HKAS126472** | **Yunnan, China** | **OR778379** |
| *A. korhinenii* | **HKAS126473** | **Yunnan, China** | **OR778378** |
| *A. korhinenii* | **WGS1417** | **Yunnan, China** | **OR778377** |
| *A. korhinenii* | WGS2045 | Yunnan, China | OR778376 |
| *A. korhinenii* | Dai25210 | Yunnan, China | OR842724 |
| *A. korhinenii* | Dai25211 | Yunnan, China | OR842725 |
| *A. korhinenii* | Dai25212 | Yunnan, China | OR842726 |
| *A. korhinenii* | Dai25213 | Yunnan, China | OR842727 |
| *A. korhinenii* | Dai25214 | Yunnan, China | OR842728 |
| *A. limonea* | CMW 4680 | New Zealand | DQ435655 |
| *A. limonea* | CMW 4991 | New Zealand | DQ435656 |
| *A. limonea* | Plim 8466 | New Zealand | FJ618654 |
| *A. luteobubalina* | CMW 4977 | Australia | DQ435657 |
| *A. luteobubalina* | CMW 8876 | Chile | DQ435658 |
| *A. mellea* | A-10 | Tokyo, Japan | AB510800 |
| *A. mellea* | AS-1 | Oita, Japan | AB539508 |
| *A. mellea* | 92-41 | Yamanashi, Japan | AB539510 |
| *A. mellea* | 97-29 | Fukushima, Japan | AB539512 |
| *A. mellea* | CMW 4613 | Iran | DQ435637 |
| *A. mellea* | ST21 | New Hampshire, USA | JF313127 |
| *A. mellea* | ST20 | Wisconsin, USA | JF313128 |
| *A. mellea* | ST5 | Virginia, USA | JF313137 |
| *A. mellea* | Am115 | Ontario, USA | JF895857 |
| *A. mellea* | FP-135350-Sp | Surrey, England | JN796025 |
| *A. mellea* | 97081/1 | Madeira, Portugal | JN796028 |
| *A. mellea* | 03277/2 | Trentino, Italy | JN796036 |
| *A. mellea* | MEX 74 | Estado de Mexico, Mexico | KC111011 |
| *A. mellea* | HKAS 85471 | Yunnan, China | KT822341 |
| *A. mellea* | HKAS 85599 | Yunnan, China | KT822342 |
| *A. mellea* | HKAS 86591 | Hebei, China | KT822343 |
| *A. mellea* | HKAS 86592 | Guizhou, China | KT822344 |
| *A. mellea* | HKAS 86593 | Zhejiang, China | KT822345 |
| *A. mellea* | HKAS 86612 | Yunnan, China | KT822346 |
| *A. mellea* | HKAS 49004 | Sichuan, China | KT822347 |
| *A. mellea* | HKAS 86598 | Africa | KT822348 |
| *A. mellea* | HKAS 86594 | Zhejiang, China | KT822349 |
| *A. mellea* | HKAS 86599 | Muguga, Kenya | KT822350 |
| *A. mellea* | HKAS 86597 | Yunnan, China | KT822351 |
| *A. mellea* | HKAS 86588 | Japan | KT822352 |
| *A. mellea* | HKAS 86611 | New Hampshire, US | KT822353 |
| *A. mellea* | HKAS 86590 | Yunnan, China | KT822354 |
| *A. mellea* | HKAS 86600 | Kenya | KT822355 |
| *A. mexicana* | MEX 85 | State of Mexico, Mexico | KR061313 |
| *A. mexicana* | MEX 87 | State of Mexico, Mexico | KR061314 |
| *A. mexicana* | MEX 88 | State of Mexico, Mexico | KR061315 |
| *A. nabsnona* | NB3 | Tottori, Japan | AB510763 |
| *A. nabsnona* | 00-16-4 | Ibaraki, Japan | AB510765 |
| *A. nabsnona* | 36586 | Aomori, Japan | AB510766 |
| *A. nabsnona* | C21 | Idaho, USA | JF313119 |
| *A. nabsnona* | M90 | British Columbia, Canada | JF313122 |
| *A. nabsnona* | ST16 | Alaska, USA | JF313124 |
| *A. nabsnona* | HKAS 85499 | Yunnan, China | KT822410 |
| *A. nabsnona* | HKAS 85523 | Yunnan, China | KT822411 |
| *A. nabsnona* | HKAS 86544 | Yunnan, China | KT822412 |
| *A. nabsnona* | HKAS 85500 | Yunnan, China | KT822413 |
| *A. novae-zelandiae* | CMW 4722 | New Zealand | DQ435652 |
| *A. novae-zelandiae* | CMW 5448 | Australia | DQ435653 |
| *A. ostoyae* | 00-10 | Mie, Japan | AB539463 |
| *A. ostoyae* | 94-72 | Iwate, Japan | AB539469 |
| *A. ostoyae* | 05-82 | Hokkaido, Japan | AB539479 |
| *A. ostoyae* | EC4 | Nurmijärvi, Finland | JF746925 |
| *A. ostoyae* | HKAS 86579 | Jilin, China | KT822428 |
| *A. ostoyae* | HKAS 86581 | Jilin, China | KT822429 |
| *A. ostoyae* | HKAS 86580 | Jilin, China | KT822430 |
| *A. ostoyae* | HKAS 86582 | Jilin, China | KT822431 |
| *A. ostoyae* | EC5 | France | JF746926 |
| *A. pallidula* | CMW 4971 | Australia | DQ435647 |
| *A. pallidula* | 3626 | Australia | FJ618665 |
| *A. puiggarii* | MCA 3111 | Guyana | KU289104 |
| *A. puiggarii* | TH 9751 | Guyana | KU289113 |
| *A. sinapina* | 35247 | Hokkaido, Japan | AB510774 |
| *A. sinapina* | 33055 | Nagano, Japan | AB539492 |
| *A. sinapina* | 2002-65 | Yamanashi, Japan | AB539493 |
| *A. sinapina* | 44702 | Hokkaido, Japan | AB539494 |
| *A. sinapina* | M50 | British Columbia, Canada | JF313114 |
| *A. sinapina* | ST13 | Michigan, USA | JF313131 |
| *A. sinapina* | ST12 | Washington, USA | JF313132 |
| *A. sinapina* | HKAS 86566 | Jilin, China | KT822422 |
| *A. sinapina* | HKAS 86567 | Jilin, China | KT822423 |
| *A. sinapina* | HKAS 86568 | Heilongjiang, China | KT822424 |
| *A. solidipes* | ST2 | Washington, USA | JF313139 |
| *A. solidipes* | P1404 | Idaho, USA | JF313140 |
| *A. solidipes* | ST1 | New Hampshire, USA | JF313141 |
| *A. xiaocaobaensis* | SWFC 12637 | Yunnan, China | MT974491 |
| *A. xiaocaobaensis* | SWFC 12638 | Yunnan, China | MT974492 |
| *A. xiaocaobaensis* | SWFC 12639 | Yunnan, China | MT974493 |
| *Armillaria* sp. 1 TG-2016 | HKAS 85519 | Yunnan, China | KT822373 |
| *Armillaria* sp. 1 TG-2016 | HKAS 85527 | Yunnan, China | KT822374 |
| *Armillaria* sp. 1 TG-2016 | HKAS 85551 | Yunnan, China | KT822382 |
| *Armillaria* sp. 1 TG-2016 | HKAS 85575 | Yunnan, China | KT822383 |
| *Armillaria* sp. 1 TG-2016 | HKAS 86615 | Guizhou, China | KT822384 |
| *Armillaria* sp. 1 TG-2016 | HKAS 85457 | Yunnan, China | KT822385 |
| *Armillaria* sp. 1 TG-2016 | HKAS 86621 | Sichuan, China | KT822386 |
| *Armillaria* sp. 1 TG-2016 | HKAS 86622 | Sichuan, China | KT822390 |
| *Armillaria* sp. 1 TG-2016 | HKAS 85581 | Yunnan, China | KT822392 |
| *Armillaria* sp. 2 TG-2016 | HKAS 86623 | Sichuan, China | KT822363 |
| *Armillaria* sp. 2 TG-2016 | HKAS 86554 | Yunnan, China | KT822366 |
| *Armillaria* sp. 2 TG-2016 | HKAS 86551 | Yunnan, China | KT822367 |
| *Armillaria* sp. 2 TG-2016 | HKAS 86557 | Yunnan, China | KT822368 |
| *Armillaria* sp. 2 TG-2016 | HKAS 86543 | Yunnan, China | KT822369 |
| *Armillaria* sp. 2 TG-2016 | HKAS 86556 | Yunnan, China | KT822370 |
| *Armillaria* sp. 3 TG-2016 | HKAS 86555 | Yunnan, China | KT822381 |
| *Armillaria* sp. 3 TG-2016 | HKAS 85449 | Yunnan, China | KT822387 |
| *Armillaria* sp. 3 TG-2016 | HKAS 86613 | Guizhou, China | KT822388 |
| *Armillaria* sp. 3 TG-2016 | HKAS 86614 | Guizhou, China | KT822391 |
| *Armillaria* sp. 3 TG-2016 | HKAS 86552 | Yunnan, China | KT822393 |
| *Armillaria* sp. 3 TG-2016 | HKAS 86553 | Yunnan, China | KT822394 |
| *Armillaria* sp. 3 TG-2016 | HKAS 86548 | Yunnan, China | KT822395 |
| *Armillaria* sp. 4 TG-2016 | HKAS 86620 | Sichuan, China | KT822389 |
| *Armillaria* sp. 5 TG-2016 | HKAS 51046 | Sichuan, China | KT822356 |
| *Armillaria* sp. 5 TG-2016 | HKAS 86601 | Heilongjiang, China | KT822357 |
| *Armillaria* sp. 5 TG-2016 | HKAS 86606 | Yunnan, China | KT822359 |
| *Armillaria* sp. 5 TG-2016 | HKAS 86619 | Hubei, China | KT822362 |
| *Armillaria* sp. 5 TG-2016 | HKAS 86610 | Xinjiang, China | KT822371 |
| *Armillaria* sp. 5 TG-2016 | HKAS 86609 | Yunnan, China | KT822372 |
| *Armillaria* sp. 5 TG-2016 | HKAS 86608 | Yunnan, China | KT822375 |
| *Armillaria* sp. 5 TG-2016 | HKAS 86618 | Hubei, China | KT822376 |
| *Armillaria* sp. 5 TG-2016 | HKAS 85594 | Yunnan, China | KT822377 |
| *Armillaria* sp. 5 TG-2016 | HKAS 86602 | Yunnan, China | KT822378 |
| *Armillaria* sp. 5 TG-2016 | HKAS 45820 | Jilin, China | KT822380 |
| *Armillaria* sp. 5 TG-2016 | HKAS 51692 | Sichuan, China | KT822415 |
| *Armillaria* sp. 6 TG-2016 | HKAS 86576 | Heilongjiang, China | KT822360 |
| *Armillaria* sp. 6 TG-2016 | HKAS 86574 | Jilin, China | KT822361 |
| *Armillaria* sp. 6 TG-2016 | HKAS 86575 | Jilin, China | KT822364 |
| *Armillaria* sp. 6 TG-2016 | HKAS 86578 | Heilongjiang, China | KT822365 |
| *Armillaria* sp. 6 TG-2016 | HKAS 86577 | Heilongjiang, China | KT822379 |
| *Armillaria* sp. 7 TG-2016 | HKAS 86607 | Yunnan, China | KT822358 |
| *Armillaria* sp. 8 TG-2016 | HKAS 86573 | Jilin, China | KT822398 |
| *Armillaria* sp. 8 TG-2016 | HKAS 86571 | Heilongjiang, China | KT822404 |
| *Armillaria* sp. 8 TG-2016 | HKAS 86560 | Shanxi, China | KT822408 |
| *Armillaria* sp. 9 TG-2016 | HKAS 45821 | Yunnan, China | KT822396 |
| *Armillaria* sp. 9 TG-2016 | HKAS 86563 | Hubei, China | KT822397 |
| *Armillaria* sp. 9 TG-2016 | HKAS 85567 | Yunnan, China | KT822399 |
| *Armillaria* sp. 9 TG-2016 | HKAS 85572 | Yunnan, China | KT822400 |
| *Armillaria* sp. 9 TG-2016 | HKAS 86561 | China | KT822401 |
| *Armillaria* sp. 9 TG-2016 | HKAS 86570 | Heilongjiang, China | KT822402 |
| *Armillaria* sp. 9 TG-2016 | HKAS 86572 | Jilin, China | KT822403 |
| *Armillaria* sp. 9 TG-2016 | HKAS 86564 | Hubei, China | KT822405 |
| *Armillaria* sp. 9 TG-2016 | HKAS 86559 | Shanxi, China | KT822406 |
| *Armillaria* sp. 9 TG-2016 | HKAS 86558 | Korea | KT822407 |
| *Armillaria* sp. 10 TG-2016 | HKAS 86541 | Yunnan, China | KT822435 |
| *Armillaria* sp. 10 TG-2016 | HKAS 83361 | Yunnan, China | KT822436 |
| *Armillaria* sp. 10 TG-2016 | HKAS 83303 | Yunnan, China | KT822437 |
| *Armillaria* sp. HKAS86549 | HKAS 86549 | Yunnan, China | KT822433 |
| *Armillaria* sp. HKAS86550 | HKAS 86550 | Yunnan, China | KT822434 |
| *Armillaria* sp. Nag.E | 96-37-1 | Kanagawa, Japan | AB510769 |
| *Armillaria* sp. Nag.E | NE4 | Tottori, Japan | AB510771 |
| *Armillaria* sp. Nag.E | 2000-23-02 | Aomori, Japan | AB510773 |
| *Desarmillaria ectypa* | Je-4 | Aomori, Japan | AB558993 |
| *D. ectypa* | Je-9 | Aomori, Japan | AB558995 |
| *D. ectypa* | BRNM704974 | Austria | EU251403 |
| *D. ectypa* | HKAS 86565 | Auvergne, France | KT822438 |
| *D. ectypa* | MY 84941 | France | FJ618643 |
| *D. ectypa* | CMW 15693 | France | FJ875698 |
| *D. tabescens* | OOI99 | USA | JF313112 |
| *D. tabescens* | ET3 | Puy de Dome, France | JF746929 |
| *D. tabescens* | 35072 | Ibaraki, Japan | AB510804 |
| *D. tabescens* | 2006-20-01 | Gunma, Japan | AB510807 |
| *D. tabescens* | 44618 | Ibaraki, Japan | AB539514 |
| *D. tabescens* | HKAS 86605 | Slovenia | KT822439 |
| *D. tabescens* | HKAS 86604 | Messina, Italy | KT822440 |
| *D. tabescens* | HKAS 86603 | Beijing, China | KT822441 |

